# Membrane compression by synaptic vesicle exocytosis triggers ultrafast endocytosis

**DOI:** 10.1101/2022.06.12.495801

**Authors:** Haoyuan Jing, Tyler Ogunmowo, Sumana Raychaudhuri, Grant F. Kusick, Yuuta Imoto, Shuo Li, Kie Itoh, Edwin R. Chapman, Taekjip Ha, Shigeki Watanabe, Jian Liu

## Abstract

Compensatory endocytosis keeps the surface area of secretory cells constant following exocytosis. At chemical synapses, clathrin-independent ultrafast endocytosis maintains such homeostasis. This endocytic pathway is temporally and spatially coupled to exocytosis, initiating within 50 ms at the region immediately next to where vesicles fuse: the active zone. How synaptic vesicle exocytosis induces ultrafast endocytosis is unknown. Here, we demonstrate that actin filaments are enriched in the region surrounding active zone at mouse hippocampal synapses and that the membrane area conservation due to this actin corral is necessary for exo-endocytic coupling. Simulations suggest that flattening of fused vesicles exerts lateral membrane pressure in the plasma membrane against the actin corral, resulting in rapid formation of endocytic pits at the border between the active zone and the surrounding actin-enriched region. Consistent with our simulations, ultrafast endocytosis does not initiate when actin organization is disrupted, either pharmacologically or by ablation of the actin-binding protein Epsin1. These data suggest that endocytosis is mechanically coupled to exocytosis at synapses.

## Main

Neurons secrete neurotransmitter through exocytosis of synaptic vesicles at chemical synapses. To sustain neurotransmission, the excess membrane added to the surface is retrieved rapidly by endocytosis. In addition to being the first step in synaptic vesicle recycling, this endocytic process restores the surface area and thereby membrane tension of the plasma membrane^1–4^. Recent studies suggest that synaptic vesicle endocytosis is clathrin-independent and occurs on a millisecond time scale^5–9^. This mechanism, ultrafast endocytosis, is temporally and spatially precise. It initiates as fast as 20-50 ms after exocytosis and always occurs at the edge of the active zone^5–7^. Furthermore, the membrane area of endocytic vesicles is roughly equal to the total surface area of synaptic vesicles exocytosed^5,6^. These factors suggest that ultrafast endocytosis is compensatory and may be mechanically coupled to exocytosis. Consistent with this notion, a recent study showed that even in the absence of calcium, synaptic vesicle exocytosis induces endocytosis^10^. In addition, previous theoretical work proposed that exocytosis could reduce membrane tension, which in turn, triggers ultrafast endocytosis^11^. However, membrane tension in neurons seems to propagate at ∼20 µm/s^12^, and therefore, is expected to equilibrate at the active zone within 10 ms of exocytosis, well before endocytic pit formation. Thus, even if membrane tension alters upon exocytosis, this model cannot account for the kinetics and location of ultrafast endocytosis.

### Actin filaments are enriched at the periphery of the active zone

Filamentous actin (F-actin) is essential for ultrafast endocytosis^5^. When neurons are treated with Latrunculin A, which sequesters G-actin monomers and perturbs F-actin formation^13,14^, ultrafast endocytosis fails completely, with no induction of membrane pit formation^5^. However, it is not clear why the actin cytoskeleton is so indispensable. To probe the role of F-actin in ultrafast endocytosis, we characterized the sub-synaptic localization of polymerized actin using stimulated emission depletion (STED) microscopy^15^. We recently discovered that a splice variant of Dynamin 1, Dyn1xA, forms condensates with Syndapin1^16^ and pre-accumulates at the ‘endocytic zone’ where ultrafast endocytosis takes place, located immediately next to the active zone (within 100 nm of the active zone edge)^17^. Thus, we used Dyn1xA as a proxy for the endocytic zone. We expressed the GFP-tagged calponin-homology domain of F-actin binding protein Utrophin (EGFP-UtrCH)^18^ in cultured mouse hippocampal neurons and visualized its localization relative to the active zone (Bassoon) and endocytic zone (Dyn1xA^17^). F-actin is present at presynaptic terminals (Fig. 1a, Extended data Fig. 1a for more example micrographs)^19,20^ and surrounds an active zone like a corral. However, it is largely excluded from the active zone (Fig. 1a,b), suggesting that the active zone contains little or no F-actin. F-actin signal peaks further from the edge of the active zone, at 100-200 nm, adjacent to Dyn1xA signals (Fig. 1a,b and Extended Data Fig. 1b). To confirm that EGFP-UtrCH probes F-actin at synapses, we treated neurons with Latrunculin A (10 µM, 1 min) to disrupt its organization or Jasplakinolide (100 nM, 2 min) to stabilize it^21,22^. After Latrunculin A treatment (Fig. 1a,b), EGFP-UtrCH signals decreased in intensity and became less abundant in regions closer to the active zone edge (Fig. 1a,b), suggesting that F-actin is disrupted at presynapses. By contrast, Jasplakinolide treatment did not affect overall abundance (Fig. 1a,b). Together, these results suggest that F-actin corrals the active zone and endocytic zone, in region hereafter referred to as the ‘periactive zone’ (Fig. 1b,c).

**Figure 1.**
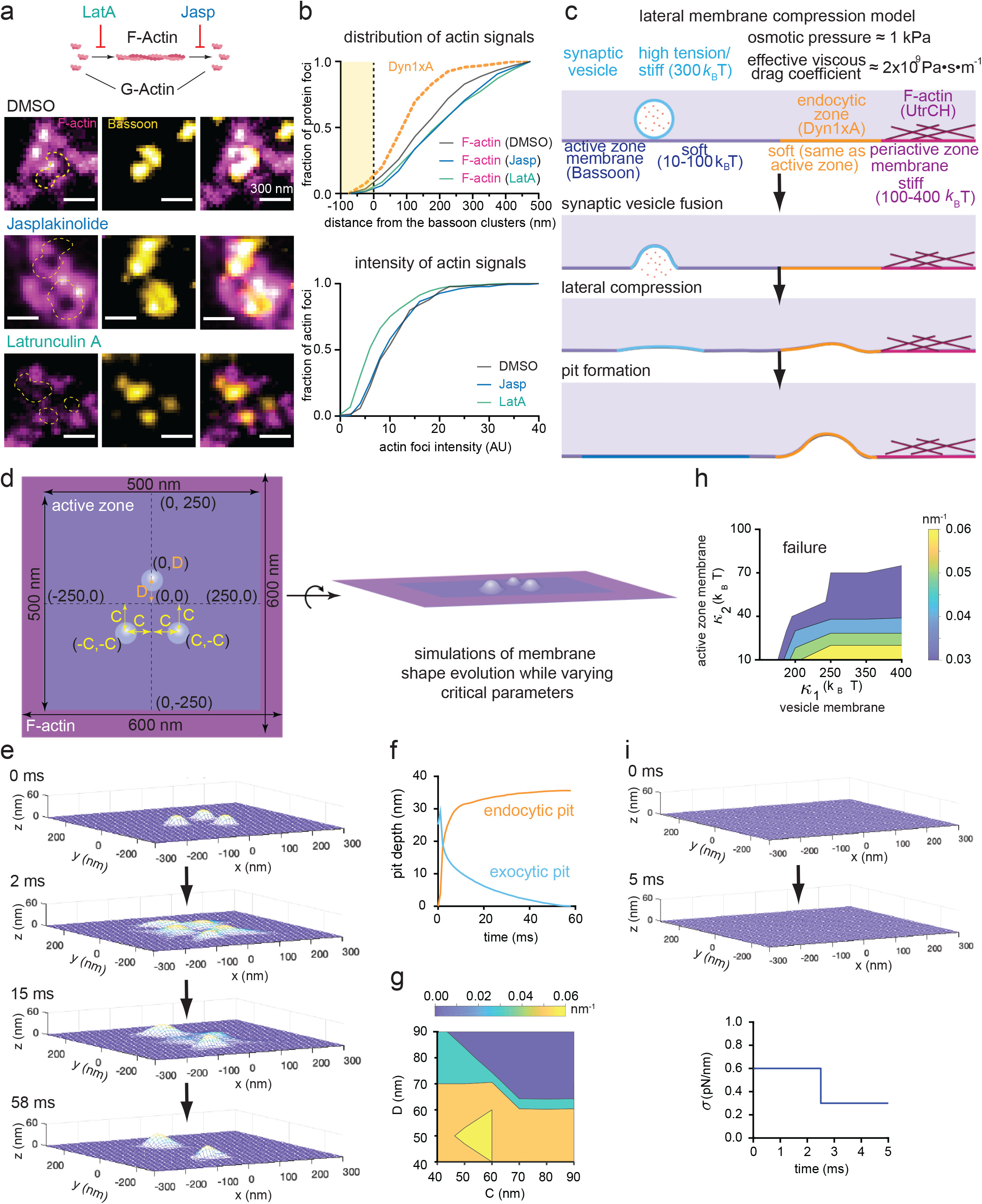
The lateral membrane compression model for ultrafast endocytosis. a. A diagram showing the effect of Latrunculin A (Lat A) and Jasplakinolide and example STED micrographs showing the localization of filamentous actin (F-actin) relative to the active zone in neurons treated with DMSO (control), Lat A, and Jasplakinolide. Active zone is marked by anti-Bassoon antibody and its secondary antibody conjugated with Alexa594. F-actin binding EGFP-UtrCH is expressed in neurons and stained with GFP-antibody and its secondary antibody conjugated with Atto646. b. Cumulative plots showing distribution of F-actin signals against the active zone boundary and intensity of F-actin signals. The active zone boundary was defined by Bassoon signals. See **Supplementary Table 2** for detailed statistical analysis. c. A schematic showing the lateral membrane compression model for ultrafast endocytosis and basic values used for simulations. The lateral membrane pressure exerted by exocytosis is predicted to compress the plasma membrane against the stiff periactive zone membrane and induce pit formation at the interface between actin-free and actin-enriched regions, or at the endocytic zone. d. Schematics showing the initial conditions of simulations from top-down view (left) and orthogonal view (right). The initial length of active zone (blue) is set at 500 nm. The width of F-actin band (purple) is set at 50 nm. Here, the active zone refers to the actin-free membrane area that includes not only the vesicle fusing area but also the endocytic zone. In contrast, the periactive zone is represented by the F-actin band where actin cortex impinges upon the membrane. The center of the active zone is set as (x, y) = (0, 0). One fusing vesicle (light blue circle) is placed at (D, 0), while two other vesicles are placed at (C, -C) and (-C, -C) such that three vesicles would form an isosceles triangle. As the initial condition, we set C = D = 60 nm. e. Snapshots from simulations, showing the evolution of membrane curvature within the active zone over time. Three fusing vesicles are organized with C = D = 60 nm. At 58 ms, simulations reach the steady state, with 2 endocytic pits forming at the boundary between active zone and actin-enriched region. f. Plot showing the depth of exocytic pits and endocytic pits as a function of time. g. Plot showing the resulting membrane curvature as a function of the spatial arrangement of fused vesicles. Distances among vesicles are modulated by changing C and D, depicted in d. h. Plot showing the dependence of successful endocytic pit formation on bending moduli of active zone and periactive zone membranes. The colored areas indicate successful formation of endocytic pits. i. Snapshots from simulations, showing the evolution of membrane shape within the active zone as a function of tension. As shown in the plot, the membrane area conservation was relaxed at 2.5 ms; consequently, the membrane tension decreases from 0.6 pN/nm to 0.3 pN/nm. This reduction in tension mimicks the expected tension change that occurs with exocytosis. Simulations reach the steady sate at 5 ms without inducing any curvature.

### A lateral membrane compression model for exo-endocytic coupling

Cortical actin stiffens the underlying plasma membrane^23–25^ and impedes the lipid flow^26–28^. Thus, the spatial arrangement of F-actin around the active zone and endocytic zone defines two mechanical properties of membranes at these locations. First, the periactive zone is stiffer than the active zone and endocytic zone. Second, the membrane area is conserved through exocytosis and endocytosis^5,29^. With these factors and the corresponding membrane mechanical parameters (Supplementary Table 1), we developed a lateral membrane compression model to explain the spatial and temporal precision of ultrafast endocytosis (Fig. 1c). This model posits that flattening of fusing vesicles squeezes the active zone membrane, causing deformation of the membrane at the interface between the actin-free (active zone) and actin-enriched regions (periactive zone; Fig. 1c). A critical assumption of this model is that the membrane of fusing vesicles remains stiffer than the active zone membrane (Fig. 1c)^30,31^. This assumption is based on our previous work that flattening of fusing synapse vesicles into the plasma membrane takes ∼10 ms ^29^ - ∼100 times longer than the time predicted for 40-nm lipid vesicles (without proteins) based on theory (0.1 ms)^32^. In addition, synaptic vesicle membranes are enriched in components such as cholesterol, sphingomyelin, and transmembrane proteins that increase the bending modulus of the membrane^33–35^. Thus, synaptic vesicle proteins and lipids are expected to largely remain within the vesicle membrane on this time scale due to their slow intrinsic diffusion (∼ 0.1 μm^2^/s or less)^36–38^, high membrane curvature at the neck of fusing vesicles, and the presence of fusion proteins at the pore^39,40^, potentially hindering diffusion. In this case, exocytic vesicle membrane would create a local stiff domain within the softer active zone membrane and compresses the plasma membrane against the corralled section of membrane. This force, in turn, causes membrane buckling at the interface between the active zone and the actin-enriched region – the site of ultrafast endocytosis^5,17^.

The model simulations start with synaptic vesicles that have fused with and nearly halfway collapsed into the plasma membrane (∼26 nm exocytic pit depth). This simulation computes the subsequent changes in the membrane shape by the steep descent of the mechanical energy of the membrane (Fig. 1d), including contributions from membrane tension, bending energy and osmotic pressure (see Supplementary Table 1). The membrane tension is treated as the Lagrange multiplier that imposes the conservation of the total membrane area. The diameter of synaptic vesicles is set at 40 nm ^5^, with a hypothetical bending modulus of 300 *k*_*B*_T (Fig. 1c). The width of the active zone is set between 100-700 nm (Fig. 1d), matching the observed parameters in mouse hippocampal synapses (average ∼300 nm in diameter)^5,29^. The bending modulus of the active zone is set at 10-100 *k*_*B*_T (Fig. 1c)^35,41,42^. The endocytic zone is assumed to have the same bending modulus as the active zone membrane and thus is included as a part of the active zone (note that no vesicles fuse in this region; Fig. 1c,d). The actin-enriched membrane patch in the periactive zone is set at the width of 100 nm with a bending modulus ranging 100-400 *k*_*B*_T and the outer boundary being pinned (Fig. 1c). In total, the presynaptic membrane area, which contains the active zone and periactive zone, spans 200-800 nm in linear dimension. Hereby, without explicitly describing endocytosis, we used the formation of resulting pits as a proxy for ultrafast endocytosis. We assumed that when the curvature of the resulting membrane pits is sufficiently large (>0.03/nm), endocytosis would process with well-defined curvature-sensing mechanisms of endocytic proteins^43,44^, some of which form condensates just outside the active zone^16,17^. From this perspective, we leveraged the model to dissect the physical mechanisms coupling exocytosis to ultrafast endocytosis at synapses.

We simulated three vesicles fusing with the active zone membrane (Fig. 1d), reflecting the average number of vesicle fusions after a single action potential at 4 mM calcium (Ca^2+^) in the external solution^29^. This calcium concentration is used in most of our previous experiments to study ultrafast endocytosis^5,7,17,45^. Our simulations show that as the vesicles flatten out, the active zone membrane starts to deform, creating lateral pressure against the perizactive zone membrane. By ∼30 ms, this deformation turns into membrane pits at the interface between the active zone and the actin-enriched edge (Fig. 1e,f; Supplementary Movie 1). The time scale of the process and the location of pit formation are consistent with those observed by ultrastructural analysis^5,6^. The resulting curvature of endocytic pits depends on locations of fusing vesicles (Fig. 1g); the further they are apart from each other, the shallower the resulting endocytic pits are. Furthermore, if synaptic vesicle membrane is too soft (below ∼230 *k*_B_T) or active zone membrane too stiff (above ∼50 *k*_B_T), endocytic pits do not form, further suggesting that the relative stiffness is key to the mechanical coupling (Fig. 1h).

Typically, our model imposes a global constraint of membrane area conservation, from which we compute the membrane tension as the model output. To directly assess the role of membrane tension in ultrafast endocytosis, we turned off the global constraint of membrane area conservation throughout the simulation and used the membrane tension as the model input. This membrane tension scales with the energy penalty for membrane area changes. As the membrane tension decreases from 0.6 pN/nm to 0.3 pN/nm at 2.5 ms (Fig. 1i; Supplementary Movie 2), mimicking the expected tension change that occurs with exocytosis, simulations reach a steady state at 5 ms without inducing any endocytic curvature. This result suggests that simply lowering the membrane tension in the active zone is not sufficient to initiate the formation of endocytic pits (Fig. 1i) and that addition of membrane is essential for pit formation, as previously demonstrated^5,6,10^. Thus, lateral membrane compression may be the basis of coupling between exocytosis and ultrafast endocytosis at synapses.

### Membrane area conservation by stable F-actin is necessary for ultrafast endocytosis

The membrane compression model predicts that the role of actin filaments in ultrafast endocytosis is passive - they are only needed to conserve membrane area, and thus stable actin polymers are sufficient. To test this prediction, we ran simulations with three fused vesicles and removed the area conservation at 2 ms, 5 ms, or 12 ms to make the bending moduli of active zone and periactive zone equal (Fig. 2a). The model predicts that the formation of pits would fail as soon as the membrane area conservation is removed (Fig. 2a,b, Supplementary Movie 3). Thus, F-actin organization around the active zone may be essential for ultrafast endocytosis^5^.

**Figure 2.**
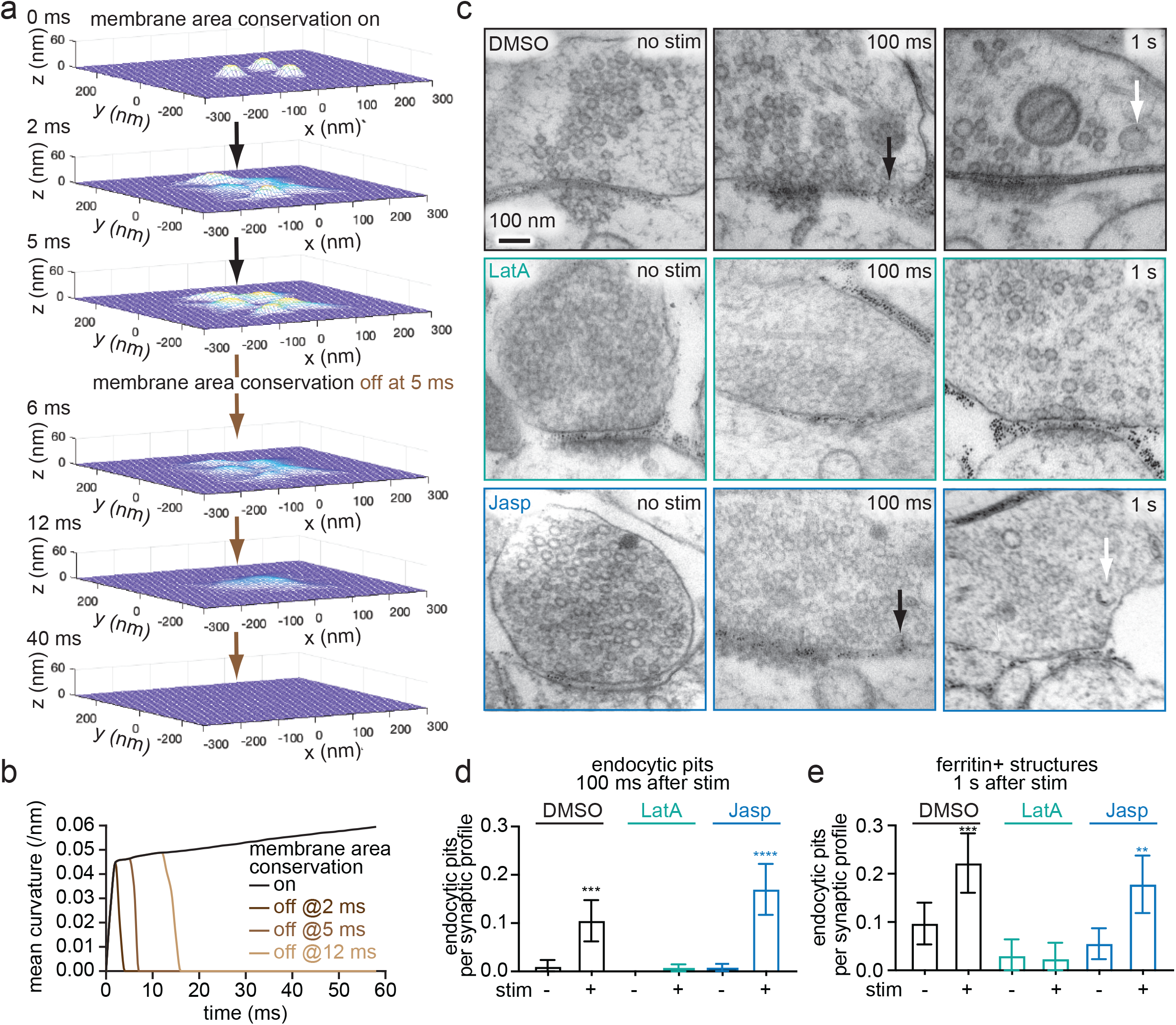
Ultrafast endocytosis is mediated by the membrane area conservation imposed by F-actin. a. Snapshots from simulations, showing the evolution of membrane curvature within the active zone by fusion of three vesicles when the membrane area conservation is initially on but turned off at 5 ms. Note that the bending moduli of the active zone and periactive zone are made uniform at 20 *k*_B_T when the membrane area conservation is turned off during simulations. b. Plot showing the temporal evolution of endocytic membrane curvature when the membrane area conservation is turned off at the indicated time points. c. Electron micrographs showing ChetaTC-expressing wild-type neurons, treated with 0.1% DMSO, 10 µM Latrunculin A (LatA), and 100 nM Jasplakinolide. The left panel show unstimulated conditions, while the right panel show 100 ms after single stimulus (10 ms light pulse, 37 °C, 4 mM external Ca^2+^). Black arrow: endocytic pit. d. Plot showing the number of endocytic pits at 100 ms after stimulation in unstimulated (-) and stimulated (+) synapses, treated with indicated drugs. e. Plot showing the number of ferritin-positive endocytic vesicles and endosomes at 1 s after stimulation in unstimulated (-) and stimulated (+) synapses, treated with indicated drugs. Mean and 95% confidential interval are shown. Kruskal-Wallis test, with Dunn’s multiple comparisons test. **p<0.01. ***p<0.001. ****p<0.0001. p values are only shown for direct comparison between unstimulated and stimulated neurons treated with the same drug. See **Supplementary Table 2** for the detailed numbers for each sample.

To experimentally test the importance of the membrane area conservation by F-actin, we reduced a total pool of F-actin with Latrunculin A (10 µM, 1 min) and performed flash-and-freeze experiments (4 mM Ca^2+^, 37 °C). As in previous experiments^5^, we applied a single 10 ms light pulse (488 nm) to neurons expressing a variant of Channelrhodopsin (ChetaTC)^46^ and froze them at 100 ms or 1 s after stimulation, with unstimulated samples serving as controls. Ferritin particles (2 mg/ml, 5 min) are applied exogenously before freezing to track recently endocytosed structures^5^. Samples were then processed for electron microscopy (see **Methods** for details). Consistent with our previous work^5^, treatment with DMSO (control) did not affect ultrafast endocytosis, while Latrunculin A treatment completely blocked it (Fig. 2c,d; more micrographs in Extended Data Fig. 2). In Latrunculin A-treated neurons, no endocytic pits formed, and consequently, ferritin-positive endocytic structures did not accumulate in synaptic terminals by 1 s (Fig. 2c,e). These data suggest that the presence, rather than the dynamics, of actin cortex underlies ultrafast endocytosis.

Endocytosis typically requires force generation by active actin polymerization at the maturing endocytic pit^47–50^. The effect of Latrunculin A could be due to the loss of such actin dynamics. By contrast, in our lateral membrane compression model only an intact cytoskeleton is required. To differentiate between these possibilities, we repeated the flash-and-freeze experiments in neurons treated with Jasplakinolide (100 nM, 2 min), which stabilizes F-actin and suppresses F-actin turnover^21^ (Fig. 1a,b). Typically, endocytosis that requires active actin polymerization is blocked by both treatments^51^. However, ultrafast endocytosis still occurred in these neurons (Fig. 2c,d). Ferritin particles were internalized into endocytic vesicles and endosomes by 1 s (Fig. 2c,e), suggesting that the initiation and kinetics of ultrafast endocytosis are unaffected by Jasplakinolide. These data suggest that a stable actin cortex, but not dynamic actin polymerization, is necessary for ultrafast endocytosis.

### Two or more vesicles must fuse to trigger ultrafast endocytosis

The lateral membrane compression model predicts that for exocytosis-induced lateral pressure to trigger endocytosis, there must be a proper scaling between the number of fusing vesicles and the dimensions of presynaptic terminals (Fig. 3a, b). If too many vesicles attempt to fuse in a small presynaptic terminal, flattening out synaptic vesicle membrane would incur a high bending energy penalty and not fully collapse. Therefore, the plasma membrane would not be pressed against the periactive zone, and endocytosis would not initiate. On the other hand, if too few vesicles fuse in a large presynaptic terminal, the lateral membrane compression would be insufficient to induce significant curvature in the endocytic zone.

**Figure 3.**
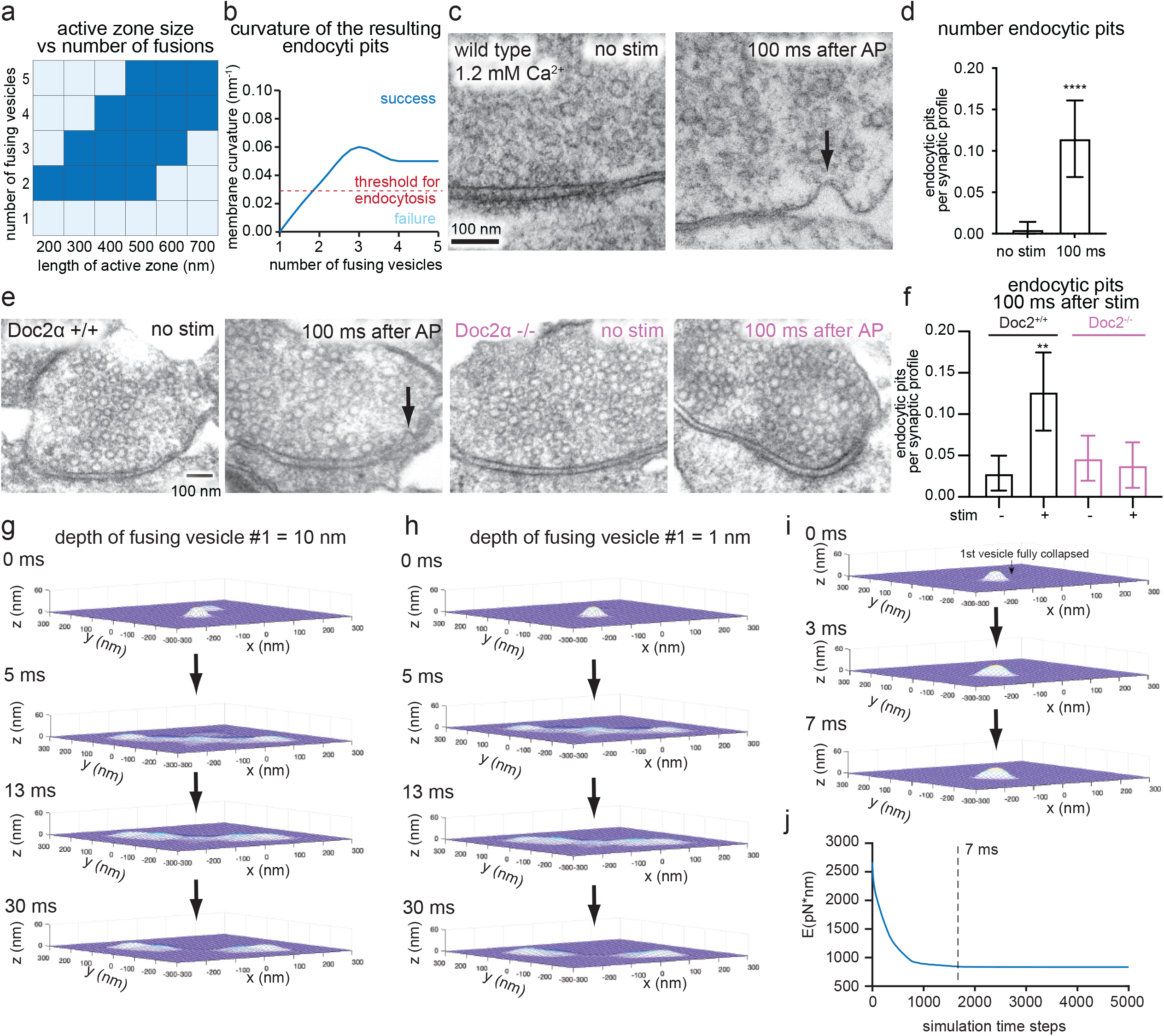
Successful exo-endocytosis coupling requires two or more vesicles to fuse simultaneously or sequentially. a. Phase diagram showing the relationship between the number fusing vesicles and dimension of the active zone for successful formation of endocytic pits (dark blue). b. Plot showing curvature of the resulting endocytic pits as a function of the number of fusing vesicles. The red dotted line indicated the threshold for endocytic proteins to recognize the curvature for further maturation of pits into vesicles. c. Example micrographs showing wild-type synapses unstimulated (left) or stimulated with electric field for 1 ms and frozen 100 ms later (right). The external calcium concentration is 1.2 mM. Black arrow: endocytic pit. d. Number of endocytic pits at 100 ms after stimulation. Mean and 95% confidential interval are shown. ****p<0.0001. Mann-Whitney test. e. Example micrographs showing wild-type and Doc2α knockout synapses unstimulated or stimulated with electric field for 1 ms and frozen 100 ms later. Black arrow: endocytic pit. f. Number of endocytic pits at 100 ms after stimulation. **p<0.01. p values are only shown for direct comparison between unstimulated and stimulated neurons of the same genotype. Brown-Forsythe and Welch ANOVA analysis, with Games-Howell multiple comparisons test. See Supplementary Table 2 for the detailed numbers for each sample. g,h. Snapshots from simulations, showing the evolution of membrane curvature within the active zone when the second vesicle is fused when the depth of first vesicle is at 10 nm (g) and 1 nm (h). i. Snapshots from simulations, showing the evolution of membrane curvature within the active zone when the second vesicle is fused much later, after the first vesicle completely flattens out by relaxing the membrane area conservation. The membrane area conservation is resumed for the second fused vesicle. j. Simulation result showing the mechanical energy evolution of the system over simulation timesteps, corresponding to (i). The energy is equilibrated within ∼ 7 ms.

Specifically, we simulated a varying number of fused vesicles while holding presynaptic terminal size constant and tested whether endocytic pits form. When we simulated two fused vesicles within typically sized active zones (200-500 nm), endocytic pits formed normally, with a curvature of 0.03 nm^-1^ (Fig. 3a, b). By contrast, when only one synaptic vesicle fuses, it cannot completely flatten out unless the membrane area conservation is transiently relaxed to restore the total membrane area of the active zone (Extended Data Fig. 3a). The lateral membrane pressure from a single vesicle is insufficient to invoke the endocytic pit formation; instead, the mechanical energy of the membrane reaches a steady-state with incomplete vesicle flattening (Extended Data Fig. 3a; curvature = 0.009 nm^-1^ at 7ms). On the other hand, when 4 vesicles fuse, endocytic pits form only if the size of the active zone is larger than 400 nm. Similarly, when 5 vesicles fuse, the size of active zone must be larger than 500 nm to accommodate extra membrane to ensure successful pit formation (Fig. 3a). Thus, the number of fusion events must be correlated with the size of presynaptic terminals^52^ for successful exo-endocytosis coupling.

These simulations predict that exo-endocytosis coupling requires at least two synaptic vesicles to fuse. To test this prediction, we performed time-resolved electron microscopy analysis of mouse hippocampal neurons using zap-and-freeze^29^. With this approach, we can deliver an electrical pulse of 1 ms to trigger a single action potential and freeze the stimulated neurons at defined time points after stimulation. Using zap-and-freeze, we previously found that fusion of multiple vesicles at a single active zone per action potential (multivesicular release) is common in cultured mouse hippocampal synapses^53,54^. However, with a physiological extracellular calcium concentration (∼1 mM)^55^, only 1-2 vesicles fuse on average from synapses that respond to a single action potential^29,53^. At this calcium concentration, 62% of responding synapses have only one vesicle fusion immediately after stimulation^29^. In contrast, at 4 mM Ca^2+^, which was used in all other experiments described here and in our previous studies, 39% of responding synapses have only one vesicle fusion. Based on our model, the frequency of ultrafast endocytosis would be expected to differ between synapses under these two calcium concentrations. However, ultrafast endocytosis occurred at a similar frequency in 1.2 mM extracellular calcium as in 4 mM calcium (Fig. 3c,d, Extended Data Fig. 4a). This is consistent with our previous study comparing 2 mM and 4 mM calcium on ultrafast endocytosis^7^. Thus, there is disparity between theoretical and experimental results.

One potential contributor to this disparity may be asynchronous fusion of synaptic vesicles. Synaptic vesicles can fuse synchronously within a few milliseconds of an action potential and asynchronously thereafter over tens and hundreds of miliseconds^56^. Based on previous studies with zap-and-freeze, patch-clamp electrophysiology measurements, and glutamate imaging, ∼35% of the total release that occurs after a single action potential is contributed by this asynchronous component in mouse hippocampal excitatory synapses^53,57^. To test whether asynchronous release contributes to ultrafast endocytosis, we performed zap-and-freeze experiments in a mutant lacking Doc2α, a calcium sensor that mediates asynchronous release after a single stimulus^57,58^. Ultrafast endocytosis failed to initiate after a single action potential in Doc2α knockouts (Fig. 3e,f), suggesting that asynchronous release helps induce ultrafast endocytosis.

In light of this observation, we next adapted the model to fuse two vesicles sequentially instead of simultaneously. The model starts with two fused vesicles: one nearly flattened out (the depth of pits: 1-10 nm) and another just starting to collapse (Fig. 3g,h, Supplementary Movies 4&5). This simulation suggests that fusion of an additional vesicle leads to the formation of membrane pits if the second vesicle fuses before the first completely flattens out (∼10 ms after an action potential^29^) (Fig. 3g,h). By contrast, when the second vesicle fuses much later, each event acts independently. As in single vesicle fusion (Extended Data Fig. 3), each fused vesicle cannot completely flatten out by itself, unless the membrane area conservation is transiently relaxed. Consequently, no endocytic pits would form, regardless of how lipids and proteins diffuse out of the first vesicle’s membrane (Fig. 3i, j). Therefore, our model suggests that asynchronous fusion likely contributes to ultrafast endocytosis under physiological, low release probability conditions. Given that synapses do not always fuse two vesicles either simultaneously or sequentially before the first completely flattens out, some exocytic events may not trigger successful ultrafast endocytic events. When ultrafast endocytosis is blocked, a slow clathrin-mediated endocytic pathway can kick in^7^ - this is consistent with observations from live-cell imaging of endocytosis^59–61^.

### Epsin1 localizes F-actin to the periactive zone

F-actin corral organization is essential to the mechanical coupling of exocytosis and endocytosis at synapses by conserving the membrane area within the active zone and endocytic zone. What actin regulators dictate F-actin organization at synapses? Epsin1 has been implicated in endocytosis at synapses^62^ and can interact with both the plasma membrane and F-actin through its N-terminal ENTH domain and C-terminal tail, respectively (Fig. 4a)^63^. If Epsin1 is important for F-actin organization, it should localize to the periphery of the active zone. To test this prediction, we determined its localization at synapses using STED microscopy^17^. Like F-actin (Fig. 1a,b), Epsin1 is localized at the periactive zone (Fig. 4b,c). To test whether Epsin1 is required for the normal enrichment of F-actin at this site, we generated an shRNA against Epsin1. Western blot analysis in mouse hippocampal cultures showed that Epsin 1 expression was reduced by 94.4 ± 2.3 % at 14 days *in vitro* (DIV) when neurons were infected with lentivirus carrying shRNA on DIV7 (Extended Data Fig. 5a). Like in neurons treated with Latrunculin A (Fig. 1a, b), Epsin1 knock-down (KD) neurons displayed 36% reduction in the F-actin level at presynaptic terminals (Fig. 4d,e). These data suggest that Epsin1 localizes F-actin to presynaptic boutons.

**Figure 4.**
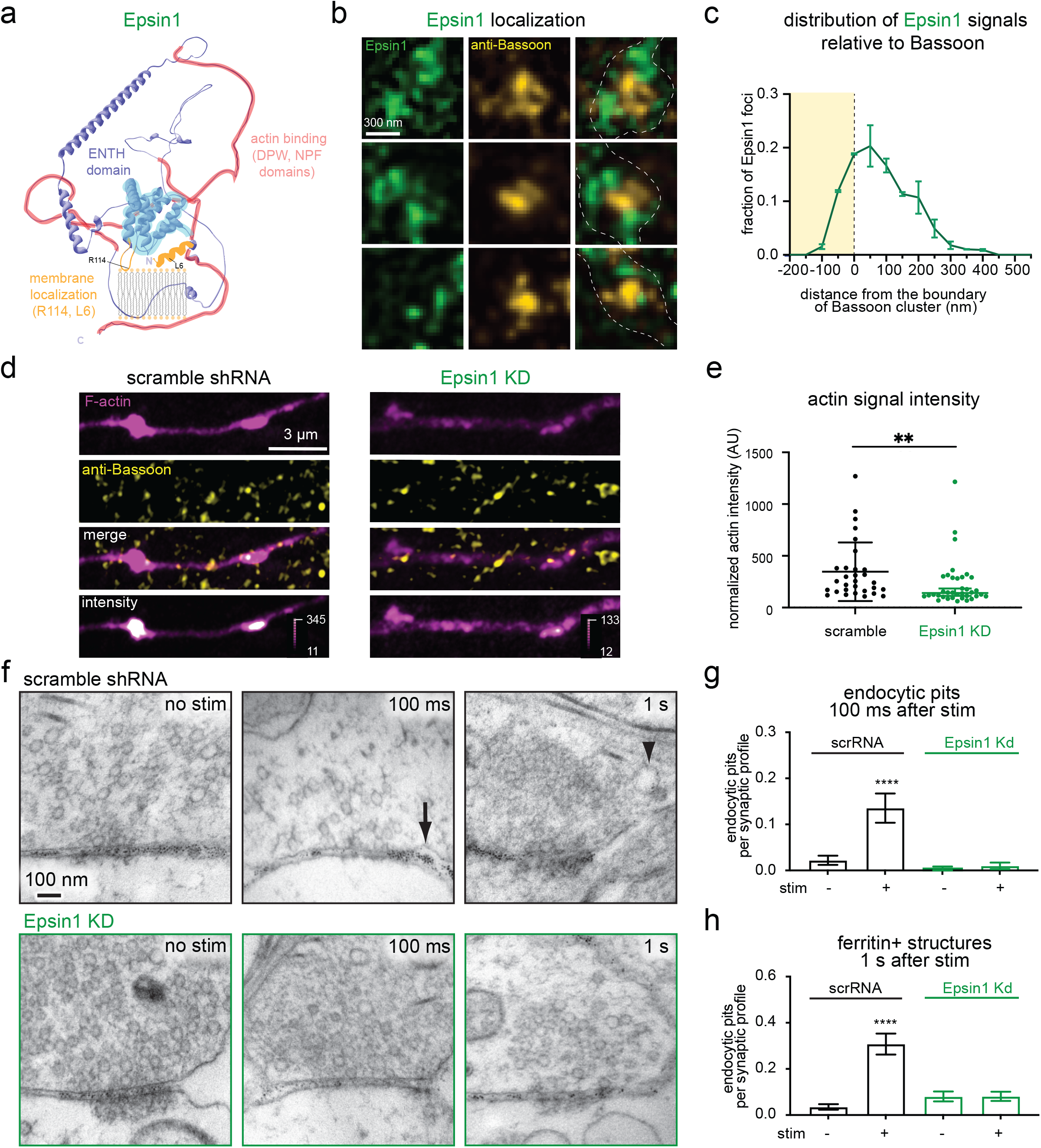
Epsin1 organizes F-actin at presynaptic terminals and is required for ultrafast endocytosis. a. A schematic showing the protein structural elements of Epsin1. Epsin1 contains ENTH domain, NPF and DPW. The C-terminal domain, marked light red, interacts with F-actin, while the ENTH domain, marked light blue, interacts with plasma membrane. b. Example STED micrographs showing the localization of Epsin 1 relative to the active zone in wild-type neurons. Active zone is marked by anti-Bassoon antibody and its secondary antibody conjugated with Alexa594. Epsin1 stained with Epsin1-antibody and its secondary antibody conjugated with Atto646. c. The distribution of Epsin1 signals against the active zone boundary. The active zone boundary was defined by Bassoon signals. See **Methods** for the analysis method. d. Example confocal fluorescence micrographs showing F-actin signals at wild-type or Epsin1 knock-down (KD) neurons. F-actin binding EGFP-UtrCH is expressed in neurons and stained with GFP-antibody and its secondary antibody conjugated with Atto646. False-colored images of the bottom panels show relative fluorescence intensity of EGFP-UtrCH. e. The normalized intensity of F-actin signals from neurons expressing scramble (scr) shRNA or Epsin1 shRNA, measured by Airyscan. Signals are normalized the fluorescence signals in axons. f. Example electron micrographs showing wild-type and Epsin1 KD synapses unstimulated or stimulated with a single electrical pulse (1 ms) and frozen 100 ms or 1 s later. Black arrow: endocytic pit. Black arrowhead: ferritin-positive endosomes. g,h. Number of endocytic pits at 100 ms after stimulation (g) or ferritin-positive structures at 1 s after stimulation (h) in neurons expressing scramble shRNA (scrRNA) or Epsin1 shRNA. Mean and 95% confidential interval are shown. Brown-Forsythe and Welch ANOVA analysis, with Games-Howell multiple comparisons test. ****p<0.0001. p values are only shown for direct comparison between unstimulated and stimulated neurons treated with the same drug. See **Supplementary Table 2** for the detailed numbers for each sample.

Our model predicts that ultrafast endocytosis would fail when actin organization is disrupted, and Latrunculin A treatment had just such an effect (Fig. 2). To test if organization of F-actin by Epsin1 is similarly required, we performed zap-and-freeze experiments in Epsin1 KD neurons (4 mM Ca^2+^, 37 °C). Knocking down Epsin1 caused defect in induction of ultrafast endocytosis, like in Latrunculin A-treated neurons. In Epsin1 KD neurons, endocytic pits failed to form within 100 ms (Fig. 4f,g, and Extended Data Fig. 5b) and no ferritin-positive endocytic vesicles and endosomes were observed at 1 s after stimulation (Fig. 4f, h). Together, these results suggest that Epsin1 organizes F-actin at the presynaptic boutons for ultrafast endocytosis.

## Discussion

We identified two essential mechanical factors for exo-endocytosis coupling at neuronal synapses. First, flattening of fusing vesicles causes lateral pressure against the actin-enriched periactive zone. Second, the actin corral enforces conservation of the total membrane area of the active zone and endocytic zone following exocytosis. This area conservation rectifies the compression, resulting in the ultrafast formation of endocytic membrane pits at the edge of the active zone. Thereafter, endocytic proteins such as Synaptojanin1, EndophilinA, Syndapin1, and Dynamin1 can further curve the membrane and generate vesicles^17,45^.

### Membrane stiffness and tension

The model assumes collapsing synaptic vesicles to be stiffer than the active zone membrane due to high protein density and enrichment of cholesterol and sphingomyelin lipids^64^. Further, the effective stiffness of fused vesicles is likely maintained by an inhibition of protein and lipid diffusion due to highly curved membrane at the neck of fused vesicles. This increased stiffness could prevent rapid collapse of vesicles. In fact, we previously showed that vesicles collapse over 10 ms^29^ instead of ∼ 0.1 ms, as predicted by theoretical modeling with pure lipid vesicles^32^. Although a reasonable assumption, measuring the membrane stiffness of synaptic vesicles, presynaptic active zones, and periactive zones is intractable by current technologies. Testing this assumption will be a priority for our future work.

Moreover, the membrane tension of fused vesicles may differ from the active zone membrane and thereby influence exo-endocytosis coupling. For instance, one may assume that synaptic vesicle fusion increases local membrane area, resulting in a spatial gradient of membrane tension, low at the site of fusion and high at the edge of active zone. A substantial spatial gradient of membrane tension is thought to persist due to attachment of cytoskeleton to the membrane^28^. This spatial gradient is expected to relax within ∼1-10 ms of vesicle fusion, since neither the active zone nor collapsing vesicles attach to cytoskeleton. This estimation suggests that the difference in the membrane tension, if it exists, may equilibrate well before the initiation of endocytic membrane pit formation (∼30-50 ms). Hence, membrane tension cannot account for the location and the timing of ultrafast endocytosis. Indeed, our model failed to reproduce endocytic membrane pit formation when membrane tension was lowered in the absence of synaptic vesicle fusion (Fig. 1h). Consistent with this simulation, recent experiments showed that membrane tension equilibrates rapidly at presynaptic terminals^12^. Taken together, these data indicate that a change in membrane tension alone, without lateral membrane compression, is insufficient to initiate ultrafast endocytosis. Nonetheless, membrane tension may influence the conservation of membrane area, thus contributing to ultrafast exo-endocytic coupling.

### Asynchronous release and its importance for endocytosis at synapses

In the lateral membrane compression model, the number of fusing vesicles, and the resulting magnitude of compressive force, is important for triggering ultrafast endocytosis. In the various sizes of active zones we simulated, two fusing vesicles are the minimal requirement. However, two vesicles do not need to fuse simultaneously and can fuse before the first one flattens out completely. In excitatory mouse hippocampal neurons, the calcium sensor Doc2α drives the majority of asynchronous release after a single action potential^57,58^. Strikingly, in the absence of this sensor, ultrafast endocytosis fails completely, suggesting that Doc2α-mediated exocytosis is a key contributor to the initiation of membrane buckling at the endocytic zone. When ultrafast endocytosis is blocked, clathrin-mediated endocytosis takes its place^7^. Thus, in synapses with fewer fusions, endocytosis still occurs, but is expected to be slow. Consistent with this notion, several previous studies suggest that endocytosis becomes slower when neurons are treated with the high-affinity calcium chelator EGTA, which abolishes asynchronous release^65–69^. It is possible that calcium directly accelerates endocytosis, as previously suggested^65–69^. However, given that calcium dynamics and calcium sensors regulate exocytosis, the role of calcium in endocytosis may need to be reassessed. Indeed, presynaptic endocytic pits reminiscent of ultrafast endocytosis form in the absence of calcium as long as exocytic fusion is induced^10^.

### F-actin organization at presynaptic terminals

The organization of F-actin is essential to the lateral membrane compression model by restricting lipid flow. We found that F-actin is arranged in a ring-like structure surrounding the active zone. These results differ from those of a recent study using single-molecule localization microscopy of presynaptic terminals formed on polystyrene microbeads. In that system, three pools of F-actin were apparent: actin rails between the active zone and vesicles, actin corrals around the terminal^20^, and in contrast to our results, an actin mesh at the active zone. The presence of F-actin in such artificially-formed active zones may reflect cytoskeletal response due to changes in cortical tension, since a bead is much stiffer than a postsynaptic terminal^70^. F-actin was rarely observed within the active zone in our experiments. Thus, active zones likely contain little or no F-actin. This spatial arrangement of F-actin at mouse hippocampal boutons is similar to presynaptic terminals of lamprey reticulospinal cord^71^ and *Drosophila* neuromuscular junctions^72^.

### The role of actin in endocytosis

The role of actin in presynaptic endocytosis has been elusive. In yeast, active force generation by actin polymerization has been long established as the mechanism by which actin supports endocytosis^73,74^. Due to the high turgor pressure across yeast plasma membranes, this force generation is essential to push the maturing vesicle inward. However, in the case of much softer animal cell membranes, the role of actin polymerization in endocytosis is controversial^50^. Genetic ablation of actin in mammalian neurons strongly perturbs endocytosis^75^. Thus, like in yeast, active actin polymerization may be important for endocytosis at synapses. Our work suggests that 1) actin is organized in a ring-like structure outside the active zone (Fig. 1a), with the endocytic zone also largely free of F-actin, and 2) intact actin filaments, but not their dynamic polymerization, are essential for the initiation of ultrafast endocytosis (Fig. 2). In the lateral membrane compression model, stable F-actin is necessary to conserve the membrane area by restricting lipid flow. However, these do not exclude the possibility that actin dynamics are necessary for ultrafast endocytosis. In fact, ultrafast endocytosis fails within 30-60 seconds of Latrunculin A treatment, indicating that the turnover of F-actin is rapid. In addition, recent work suggests that the actin nucleator formin is required for synaptic vesicle endocytosis at small hippocampal boutons^8^. Therefore, dynamic actin polymerization may be important for maintaining the cortical F-actin organization in the periactive zone.

### Mechanisms of F-actin organization at synapses

Epsin1 seems to play a key role in actin organization at synapses. Specifically, knocking down Epsin1 in mouse hippocampal neurons results in 1) a reduced amount of F-actin around the periactive zone and 2) failure of ultrafast endocytosis. We interpret these results to indicate that Epsin1 is the organizer of cortical F-actin at synapses for ultrafast endocytosis. However, it is not clear whether Epsin1 is the sole organizer of F-actin at synapses, since it interacts with other endocytic proteins such as Eps15 and Hip1R^63,76^, both of which interact either directly or indirectly with membranes and actin. Thus, Epsin1-interacting proteins may also have roles in actin organization. In addition, the first alpha helix (α0) of the N-terminal ENTH domain can insert into the membrane and induce curvature formation (Fig 4a)^77,78^ – this function of Epsin1 is thought to be important for endocytosis. Thus, Epsin1 may have a more active role in ultrafast endocytosis beyond organizing F-actin. Nonetheless, our results here suggest that the first step in endocytosis at synapses does not involve proteins, but rather is initiated by an excess of ‘soft’ membrane in an active zone corral.

## Acknowledgements

We thank Sydney Brown and people in the Johns Hopkins Microscopy Facility for technical assistance. We thank Erik Jorgensen, Thien Vu, and Edward Hujber for active discussion. We thank Mike Seibert for developing the Doc2α genotyping PCR protocol and Ye Ma for the analysis code of STED images. S.W. was supported by start-up funds from the Johns Hopkins University School of Medicine, Johns Hopkins Discovery funds, Johns Hopkins Catalyst award, the National Science Foundation (1727260), and the National Institutes of Health (NIH; 1DP2 NS111133-01 and 1R01 NS105810-01A1) awarded to S.W. S.W. is an Alfred P. Sloan fellow, a McKnight Foundation Scholar and a Klingenstein and Simons Foundation scholar. E.R.C. was supported by the NIH (R35NS097362 and R01MH061876). Y.I. was supported by JSPS. T.O. and G.F.K. were supported by the National Science Foundation GRFP (T.O.: 2019241734, GFK: 2016217537) and a grant from the National Institutes of Health to the BCMB program of the Johns Hopkins University School of Medicine (T32 GM007445). G.F.K. was additionally supported by the Hay Fellowship from the Department of Cell Biology at Johns Hopkins University. T.H. and E.R.C. are investigators of the Howard Hughes Medical Institute. J.L was supported by start-up funds from the Johns Hopkins University School of Medicine, Johns Hopkins Catalyst award, the National Science Foundation (2105837 and 2148534).

## Author contributions

S.W. and J.L. oversaw the research. H.J., T.O., S.R., G.F.K., S.W., and J.L. conceived the study and designed the experiments. H.J. and J.L. performed theoretical work. T.O., S.R., S.L., G.F.K., Y.I. S.W. performed wet-bench work. K.I. prepared cultured neurons. T.H. provided the STED microscopy. E.R.C. provided Doc2 knockout animals. H.J., T.O., S.R., S.W., and J.L. wrote the manuscript. All authors contributed to editing. S.W. and J.L. funded research.

## Competing interests

The authors declare no competing interests.

## Methods

### Animal use

All the animal work was performed according to the National Institutes of Health guidelines for animal research with approval from the Animal Care and Use Committees at the Johns Hopkins University School of Medicine. For Doc2α knockout experiments, C57BL/6-*DOC2a*^*em1Smoc* 79^ were maintained as heterozygotes neurons culture from homozygous null P0 pups, with homozygous WT littermates used as controls. For all other experiments, neurons were cultured from E18 embryos from C57BL/6J mice.

### Primary neuron culture

Primary hippocampal neurons were isolated from either E18 embryos or P0 pups of both genders. The brains were harvested from animals and hippocampi were dissected under a binocular microscope. Dissected hippocampi were collected in ice-cold dissecting media (1 x HBSS, 1 mM sodium pyruvate, 10 mM HEPES, pH7.2-7.5, 30 mM glucose, 1% penicillin-streptomycin) and later digested with papain (0.5 mg/ml) and DNase (0.01%) for 25 min at 37°C. Cells were dissociated by trituration using fire-polished Pasteur pipettes.

For high pressure freezing experiments neurons were plated onto 6-mm sapphire disks (Technotrade Inc) coated with poly-D-lysine (1 mg/ml) and collagen (0.6 mg/ml) with an astrocyte feeder layer on it (Ref: JOVE). Cortices were harvested from E18/P0 animals and astrocyte was isolated with a treatment of trypsin (0.05%) for 20 min at 37 °C, followed by trituration. Astrocytes were seeded in T-75 flasks containing DMEM supplemented with 10% FBS and 0.2% penicillin-streptomycin. After 2 weeks, astrocytes (50K) were plated on sapphire disks at a density. After 1 week in culture, astrocytes were incubated with 5-Fluoro-2′-deoxyuridine (81 µM) and uridine (204 µM) for at least 2 hours to stop mitosis. Prior to addition of hippocampal neurons, medium was changed to Neurobasal-A (Gibco) supplemented with 2 mM GlutaMax, 2% B27 and 0.2% penicillin-streptomycin. For fluorescence imaging, dissociated hippocampal neurons were seeded onto 18-mm or 25-mm coverslips (Carolina Biologicals) coated with poly-L-lysine (1 mg/ml, Sigma) at a density of 25-40 × 10^3^ cells/cm^2^ in Neurobasal media (Gibco) supplemented with 2 mM GlutaMax, 2% B27, 5% horse serum and 1% penicillin-streptomycin (NM5) at 37 °C in 5% CO_2_. Next day, media were changed to Neurobasal media with 2 mM GlutaMax and 2% B27 (NM0), and neurons were maintained in this medium until use. For biochemical experiments, dissociated hippocampal neurons were seeded on poly-L-lysine (1mg/ml) coated plates with Neurobasal media supplemented with 2 mM GlutaMax, 2% B27, 5% horse serum and 1% penicillin-streptomycin, at a density of 1 × 10^5^ cells/cm^2^. Next day, the medium was changed to Neurobasal medium containing 2 mM GlutaMax and 2% B27 (NM0), and neurons were maintained in this medium. A half of the media was refreshed every week or as needed. For Doc2a KO and WT littermate cultures, tail clips were obtained from live P0 pups and genotyped as described^53^. Brain tissues were harvested from correct genotypes and hippocampal neurons were prepared as described above.

### Plasmids

For knocking down Epsin1, a plasmid (pLKO.1-puro backbone) containing shRNA sequence (CCGGGATGAAGAATATCGTCCACAACTCGAGTTGTGGACGATATTCTTCATCTTTTTG) was purchased from Sigma (Mission shRNA, clone ID TRCN0000111703). For non-targeting scramble control, oligo’s containing sequence (GATCCCTTCGCACCCTACTTCGTGGttcaagagaCCACGAAGTAGGGTGCGAATTTTTGGAAATTAAT) was cloned under U6 promoter. Annealed oligo’s were inserted into BamH1 and PacI sites of modified pFUGW vector using TAKARA solution I. For labeling actin, we acquired a plasmid containing tandem calponin homology (CH) domains from the human actin binding protein utrophin which are C-terminally linked to GFP from addgene (pEGFP-C1 Utr261-EGFP, #58471). To label the endocytic zone, we used a plasmid expressing Dynamin1xA C-terminally tagged to GFP, purchased from addgene (phsDyn1xA-EGFP-N1, #120313).

### Lentivirus preparation

Lentivirus containing either Epsin1 shRNA or nontargeting scramble shRNA was prepared as described earlier^17^. Briefly, shRNA construct along with two helper DNA constructs (pHR-CMV8.2 deltaR (Addgene 8454) and pCMV-VSVG (Addgene 8455)) at a 4:3:2 molar ratio was transfected into HEK293T cells using polyethylene amine. Cell supernatant containing the virus was collected 3 days after transfection and 20-fold concentrated using Amicon Ultra 15 10K (Millipore) centrifugal filter. Aliquots were flash frozen in liquid nitrogen and stored in -80 °C until use.

### Transient transfection of neurons

For transient expression of proteins, neurons were transfected at DIV (days in vitro) 14-16 by Lipofectamine 2000 (Invitrogen) according to the manufacturer’s manual. Prior to transfection, half of the media from each well was taken out and mixed with fresh NM0 (see above) that was left to warm to 37 C° and equilibrate with CO_2_ in an incubator (recovery media). The rest of the media was aspirated and replaced with fresh NM0 for transfection. Plasmids were diluted in NeuroBasal Plus (Gibco) media so that 0.5-1 µg of DNA would be added to each well. Prior to addition, DNA was mixed with a solution containing Lipofectamine 2000 such that there was a 1:1 ratio of µg DNA to µL Lipofectamine. This mixture was added to each well and incubated for 4 hours. Afterwards, the transfection media was removed and replaced with recovery media previously prepared. After 16-20 hours, neurons were either used for pharmacological treatment or fixed for immunofluorescence.

### Pharmacology

For Actin perturbation experiments, transfected cells were treated immediately prior to fixation. Latrunculin A and Jasplakinolide stock solutions were prepared in DMSO as 10 mM and 100 µM, respectively. Stock drug solutions were diluted into the media in wells at a 1:1000 dilution for a final concentration of 10 µM for Latrunculin A and 100 nM for Jasplakinolide. Cells were treated with Latrunculin A for 30 seconds to 1 minute, while cells were treated with Jasplakinolide for 1.5-2 minutes. Negative control samples were treated with 0.1% DMSO.

### Immunofluorescence

For immunofluorescence experiments were performed with DIV 14-16 hippocampal neurons. Culture media was removed from the wells and fixed with a prewarmed 1X PBS containing 4% paraformaldehyde and 4% sucrose in for 20 minutes at room temperature. Following fixation, cells were washed three times with 1x PBS. Next, cells were permeabilized by 0.2% Triton X-100 diluted in 1X PBS for 8 minutes. After three washes with 1X PBS, cells were blocked by 1% BSA in 1X PBS for 1 hour. Then, the coverslips containing neurons were transferred to a humidified chamber and placed face down on a drop of primary antibody solution. Primary antibodies were diluted 1:500 in a 1% BSA 1X PBS and cells were incubated at 4 °C overnight. Utr261-EGFP was stained by an anti-GFP rabbit polyclonal antibody (MBL International). Endogenous Bassoon protein was stained by an anti-bassoon mouse monoclonal (Synaptic Systems) antibody. Next, coverslips were washed with 1X PBS three times. Secondary antibodies were diluted in 1X PBS containing 1% BSA. For superresolution STED imaging, an anti-rabbit Atto-647N (Rockland) secondary antibody was used at 1:120 dilution and an anti-mouse Alexa 594 (Invitrogen) secondary antibody was used at 1:1000. Secondary antibody incubation was performed in a humidified chamber as described previously for 1 hour at room temperature. Following three 1X PBS washes, cells were rinsed with Milli-Q water and mounted on a glass slide containing a drop of ProLong Diamond Antifade Mounting media (Thermo Fisher). Mounting Media was allowed to solidify for 24 hours at room temperature in the dark before proceeding to STED imaging. For confocal imaging in Airyscan, cells were prepared using the same protocol.

### Stimulated emission depletion microscopy (STED)

All STED images were obtained using a home-built two-color STED microscope ^80^. A femtosecond laser beam with a repetition rate of 80 MHz from a Ti:Sapphire laser head (Mai Tai HP, Spectra-Physics) is split into two parts: one part is used to produce the excitation beam, which is coupled into a photonic crystal fiber (Newport) for wide-spectrum light generation and is further filtered by a frequency-tunable acoustic optical tunable filter (AA Opto-Electronic) for multi-color excitation. The other part of the laser pulse is temporally stretched to ∼300 ps (with two 15-cm-long glass rods and a 100-m long polarization-maintaining single-mode fiber, OZ optics), collimated, expanded, and wave-front modulated with a vortex phase plate (VPP-1, RPC photonics) for hollow STED spot generation to de-excite the fluorophores at the periphery of the excitation focus, thus improving the lateral resolution. The STED beam is set at 765 nm with power of 120 mW at the back focal plane of the objective (NA=1.4 HCX PL APO 100×, Leica), and the excitation wavelengths are set as 594 nm and 650 nm for imaging Alexa-594 and Atto-647N labeled targets, respectively. The fluorescent photons are detected by two avalanche photodiodes (SPCM-AQR-14-FC, Perkin Elmer). The images are obtained by scanning a piezo-controlled stage (Max311D, Thorlabs) controlled by the Imspector data acquisition program.

### Data analysis of STED images

A custom MATLAB code package was used to analyze actin and endocytic protein distribution relative to the active zone marked by Bassoon in STED images (Imoto et al). First, STED images were blurred with a Gaussian filter with radius of 1.2 pixels to reduce the Poisson noise, and then deconvoluted twice using the built-in deconvblind function: the first point spread function (PSF) input is measured from nonspecific antibody signal in the STED images, and the second PSF input is chosen as the returned PSF from the first run of blind deconvolution ^81^. Each time, 10 iterations are performed. Presynaptic boutons in each deconvoluted image were selected within 30×30-pixel (0.81 mm^2^) ROIs based on the varicosity shape and bassoon signal. The active zone boundary was identified as the contour that represents half of the intensity of each local intensity peak in the Bassoon channel, and the actin or endocytic protein clusters are picked as local maxima. The distances between the protein cluster centers and the active zone boundary are automatically calculated correspondingly. Actin and endocytic protein clusters over crossing the ROIs, and the Bassoon signals outside of the transfected neurons were excluded from the analysis. The MATLAB scripts are available by request.

### Airyscan imaging and data analysis

For Airyscan imaging, samples were imaged in Zeiss LSM880 (Carl Zeiss) in Airyscan mode. Fluorescence was acquired using a 63x objective lens (NA = 0.55) at 2048×2048 pixel resolution with the following settings: pixel Dwell 1.02 ms and pin hole size above the lower limit for Airyscan imaging, as computed by ZEN software (Zen Gray, 2.31 SP1). For experiments comparing fluorescence intensities between scramble and shRNA treated cells, staining and microscope settings were remained constant. Utr-EGFP was used to stain actin as described before and GFP signal intensity was used to determine actin level. Presynaptic regions were determined with Bassoon-Alexa594 signals. Axons were distinguished from dendritic processes based on their morphology, thin and lacking spines. Z-sections were taken for each presynaptic bouton along axons. Utr-EGFP intensities, depicting actin level in bouton, was quantitated in ImageJ. Bassoon-Alexa594 signal was used to determine the ROI’s, and all Utr-EGFP signals within ROIs were measured as the total signals at each synapse. Fluorescence intensity was normalized to the average signal within the corresponding cell body of each axon.

### High pressure freezing

Hippocampal neurons cultured on sapphire disks were frozen using high pressure freezer (EM ICE, Leica Microsystems). For actin dynamics perturbation, neurons were treated with Latrunculin A (10 µM for 1 minute), Jasplakinolide (100 nM for 2 minutes) and DMSO as control. Epsin1 and scramble shRNA’s were added to cells on DIV7 and freezing experiments were performed on DIV14. Ferritin (2 mg/ml) was used as a fluid-phase marker and added to the cells for 5 minutes prior to freezing. Cells were frozen in the physiological saline solution (140 mM NaCl, 2.4 mM KCl, 10 mM HEPES, 10 mM glucose; pH adjusted to 7.3 with NaOH, 300 mOsm) containing NBQX (3 µM, Tocris) and bicuculine (30 µM; Tocris), which were added to block recurrent synaptic activity. CaCl_2_ and MgCl_2_ concentrations were adjusted as needed for experiments (mentioned in results section). Zap-and-freeze experiments were performed as described (Kusick et al, 2020). After freezing, samples were transferred under liquid nitrogen to an automated freeze substitution unit, which was set at -90 °C (EM AFS2, Leica Microsystems). Using pre-cooled tweezers, samples were quickly transferred to anhydrous acetone at -90 °C. After disassembling the freezing apparatus, sapphire disks with cells were quickly moved to cryovials containing freeze substitution solutions and left inside EM AFS2. For EM experiments described in Figure 2 and Figure 4 freeze substitution was performed in solutions containing 1% glutaraldehyde, 1% osmium tetroxide, and 1% water in anhydrous acetone, which had been stored under liquid nitrogen then moved to the AFS2 immediately before use. The freeze substitution program was as follows: -90 °C for 6-10 hr (adjusted so substitution would finish in the morning), 5 °C h^-1^ to -20 °C, 12 h at -20 °C, and 10 °C h^-1^ to 20 °C. For EM experiments using neurons from Doc2a KO (Figure 3) a different freeze-substitution protocol was used that generated more consistent results. In this protocol, after freezing samples were first left in 0.1% tannic acid and 1% glutaraldehyde at -90 °C for ∼36 hours, then washed 5 times, once every 30 min, with pre-chilled acetone, and transferred to 2% osmium tetroxide in acetone. Freeze substitution program was as follows: 11 hr at -90 °C, 5 °C h^-1^ to -20 °C, -20 °C for 12 hr, 10 °C h^-1^ to 4 °C, then removed from the freeze substitution chamber and warmed at room temperature for ∼15 min before infiltration and embedding. For this latter protocol, all the steps were performed in universal sample containers (Leica Microsystems) and kept covered in aclar film to prevent evaporation.

### Sample preparation for electron microscopy

Following freeze-substitution, fixatives were washed with anhydrous acetone for five times, 10 min each. 100% epon araldite (epon 6.2 g; araldite 4.4 g; DDSA 12.2 g, and BDMA 0.8 ml) solution was prepared and diluted acetone to get 30%, 70% and 90% solutions. Samples were infiltrated for at least two hours at room temperature sequentially in 30%, 70% epon-araldite. Samples were then transferred to caps of polyethylene BEEM capsules with 90 % epon araldite and incubate overnight at 4 °C. Next day, samples were transferred to new caps with fresh 100 % epon araldite, changed every 2 hours three times, after which samples were cured at 60 °C for 48 hours.

After resin was cured, 40 nm sections were cut using an ultramicrotome (EM UC7, Leica microsystems) and collected on single-slot copper grids coated with 0.7 % pioloform. The sections were stained with 2.5 % uranyl acetate in 75 % methanol.

### Electron microscopy imaging and data analysis

Samples were imaged on a Hitachi 7600 TEM equipped with an AMT XR50 camera run on AMT Capture v6 (pixel size = 560 pm), at 80 kV on the 100,000x setting. Samples were blinded before imaging. Synapses were identified by a vesicle-filled presynaptic bouton and a postsynaptic density. Postsynaptic densities are often subtle in our samples, but synaptic clefts were also identifiable by 1) their characteristic width, 2) the apposed membranes following each other closely, and 3) vesicles near the presynaptic active zone. 125-150 micrographs per sample of anything that appeared to be a synapse were taken without close examination. All images were from different synapses.

EM image analysis was performed as previously described ^29^. All the images from a single experiment were randomized for analysis as a single pool. Only after this randomization were any images excluded from analysis, either because they appeared to not contain a bona fide synapse or the morphology was too poor for reliable annotation. The plasma membrane, the active zone, exocytic and endocytic pits, clathrin coated pits docked SVs, and all SVs in the bouton were annotated in ImageJ using SynapsEM plugins [https://github.com/shigekiwatanabe/SynapsEM copy archived at <u>swh:1:rev:11a6227cd5951bf5e077cb9b3220553b506eadbe</u>] (Watanabe et al 2020). To minimize bias and error, and to maintain consistency, all image segmentation, still in the form of randomized files, was thoroughly checked and edited by a second member of the lab. Features were then quantitated using the SynapsEM (Watanabe et al., 2020) family of MATLAB (MathWorks) scripts (https://github.com/shigekiwatanabe/SynapsEM). Example electron micrographs shown were adjusted in brightness and contrast to different degrees (depending on the varying brightness and contrast of the raw images), rotated, and cropped in ImageJ before import into Adobe Illustrator.

### Western blot analysis

To test efficiency of Epsin1 shRNA to knock down endogenous protein, lentivirus containing scramble or shRNA was added to cultured hippocampal neurons on DIV7. Neurons were harvested on DIV14 and lysed by addition of lysis buffer (50 mM Tris pH 8.0 and 1% SDS containing cOmplete Mini Protease Inhibitor (Roche)) and boiling at 95°C for 5 min. Lysates were centrifuged at 15,000 x g for 15 min at 4 °C and the supernatants were separated in SDS–PAGE and transferred onto Immobilon-FL membranes (Millipore). Following blocking with 5% skim milk in PBS containing 0.05% Tween-20 (PBST) for 30 min, membranes were incubated with Epsin 1(Thermo Fisher) and control β-actin (SYSY) antibodies diluted in 3% BSA/PBST overnight at 4°C, followed by IRDye secondary antibodies (LiCor) diluted in 1:10,000 in 3% BSA/PBST for 45 min at room temperature. Signal was detected using LiCor Odyssey Clx and quantification was done by Image Studio Lite from LiCor.

### Statistical analysis

Detailed statistical information is collated in table S2. STED images were acquired from ∼2 biological replicates per condition. Each replicate was a dissociated mouse hippocampal culture (N) taken from different mice on different days. For each N, roughly 30 presynaptic bouton regions of interest (ROIs) (n) were imaged from multiple transfected cells. ROIs from each replicate were pooled and quantified as previously described^16^. An alpha of 0.05 was set for null hypothesis testing. For actin and dynamin1xA foci distance statistical analysis, pooled distance measurements from each condition were assessed for distribution normality. A full-pairwise Kruskal Wallis test was performed. Afterwards, each condition was compared by Dunn’s multiple comparisons test. Intensity measurements taken from the same datasets were assessed for normality. Following this, each condition was compared by a Kruskal-Wallis test for non-parametric distributions, followed by a Dunn’s multiple comparisons test. Non-STED fluorescence experiments were prepared and quantified in the same manner as STED experiments. However, these datasets only required comparison of only two conditions, and therefore, a Mann-Whitney test was used for nonparametric statistical comparison.

For electron microscopy data, measurements were taken from roughly 100 synaptic profile micrographs (n) per condition. Replicate high-pressure freezing experiments (N) were conducted with cultures taken from different mice on different days. Sample sizes for each replicate were inferred from previous flash-and-freeze experiments as opposed to power analysis. An alpha of 0.05 was set for null hypothesis testing. For count data sets such as these, anormal, nonparametric distributions are assumed and typical. However, means are best to represent central tendency, and these data are binomially distributed. So, an ANOVA test with a Brown-Forsythe correction and Games-Howell post hoc was conducted. In the case of electron microscopy data sets with measurements of 0, Brown-Forsythe correction fails. Therefore, statistical comparison by a Kruskal-Wallis test followed by Dunn’s multiple comparisons was used instead.

### Model simulations

The mechanical energy of the membrane deformation is described by Eq. [1], using general form of Helfrich-like energy^82^:

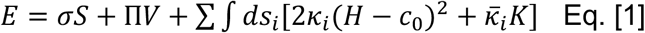

Here, *σ* is the Lagrange multiplier to enforce surface area conservation and has a unit of surface tension. *S* is the total area of the simulated membrane patch, including the membrane from the fusing synaptic vesicle, the active zone, the endocytic zone, and the peri-active zone. Π is osmotic pressure in neurons, pointing outward. *V* is the volume sandwiched between the membrane shape deformation and the flat baseline; with this, when the membrane curve inward, it causes the energy penalty from the osmotic pressure. *κ*_*i*_ is the bending modulus of the corresponding membrane, and 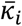 the Gaussian bending modulus. *H* = (*c*_1_ + *c*_2_)/2 is the mean curvature, and *K* = *c*_1_*c*_2_ is the Gaussian curvature, where *c*_1_ and *c*_2_ are the curvatures in the two principle directions along the two-dimensional manifold of membrane surface, respectively. *c*_0_ is the spontaneous curvature of the membrane and assumed to be zero. Since our model does not describe the initiation of exocytosis and completion of endocytosis, the topology of the membrane patch remains constant and hence, the contribution of the Gaussian curvature is constant and can be omitted^83^.

Taken together, we simplified the above equation as follows:

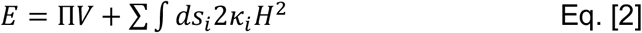

with the global constraint that the total membrane area is conserved.

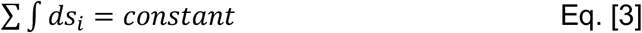

In our simulations, the membrane patch can be divided into 3 different sub-domains with different bending modulus. The first sub-domain is fusing vesicles with a bending modulus of about 100∼400 *k*_*B*_*T*. The second sub-domain includes active zone and endocytic zone and has a bending modulus of about 10∼100*k*_*B*_*T*. The last sub-domain is the actin-enriched periactive zone, surrounding the second sub-domain. The bending modulus of this domain is set at 100∼400 *k*_*B*_*T*.

The time evolution of the membrane shape is determined by the minimization of the Helfrich free energy by descending gradient method via a free software Surface Evolver^84^, which is specialized in minimizing the free energy of triangulated surfaces^85,86^. There are three constraints implemented in the simulation: 1) The total membrane area was conserved for a period of duration, which can be modulated to mimic the overall effect of this conservation; 2) the outer boundary of the actin-enriched domain was pinned to keep the membrane anchored; and 3) only the membrane shape change towards the inside of the cell is allowed. Additionally, we used measured viscous drag coefficient of the membrane shape change to scale the simulation timestep, from which we determined the real timescale of the dynamic process. Further, we leveraged adaptive timestep to simulate the membrane shape change so that the simulation timestep will become smaller and smaller as the system evolves to a steady state, where the gradient of energy landscape approaches to zero.

## Code availability

Custom R, Matlab and Fiji scripts for electron microscopy analysis are available at https://github.com/shigekiwatanabe/SynapsEM. Modeling scripts are available at https://github.com/jliu187/membrane-compression-model. More details can be found in ^87^.

## Data availability

Originals images used in this work will be uploaded to Figshare. Data will be available upon request.

---

**Supplementary Table 1:** The nominal parameters used in simulations.

| **Model Parameter** | **Physical Range** | **Nominal Value used in the model** |
| --- | --- | --- |
| Bending modulus of post-fusion vesicle, $\kappa_1$ | $\sim 10^2 k_B T$ <sup>33–35,42</sup> | $300 k_B T$ |
| Bending modulus of the actin-free membrane domains (including the active zone and endocytic zone), $\kappa_2$ | $\sim 10^1 k_B T$ <sup>35,41,42</sup> | $20 k_B T$ |
| Bending modulus of the actin-enriched membrane domain, $\kappa_3$ | $\sim 10^2 k_B T$ <sup>23–25</sup> | $300 k_B T$ |
| Osmotic pressure | $\sim 1 \text{ kPa}$ <sup>88</sup> | $1 \text{ kPa}$ |
| Viscous drag coefficient for membrane shape change | $\sim 2 \times 10^9 \text{ Pa} \cdot \text{s} \cdot \text{m}^{-1}$ <sup>89</sup> | $2 \times 10^9 \text{ Pa} \cdot \text{s} \cdot \text{m}^{-1}$ |

## Notes

### Competing Interest Statement

The authors have declared no competing interest.

