## Supplementary Figures for "Membrane compression by synaptic vesicle exocytosis triggers ultrafast endocytosis"

Extended Data Figure 1

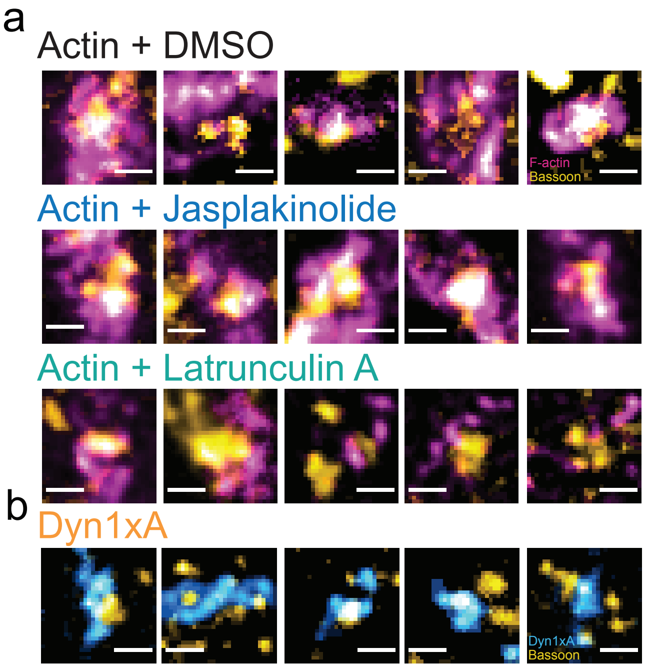

**Extended Data Figure 1.**

a. Additional example STED micrographs showing the localization of filamentous actin (F-actin) relative to the active zone in neurons treated with DMSO (control), Latrunculin A (Lat A), and Jasplakinolide. Active zone is marked by anti-Bassoon antibody and its secondary antibody.

b. Example STED micrographs showing the localization of Dynamin1xA (Dyn1xA) relative to the active zone.

Extended Data Figure 2

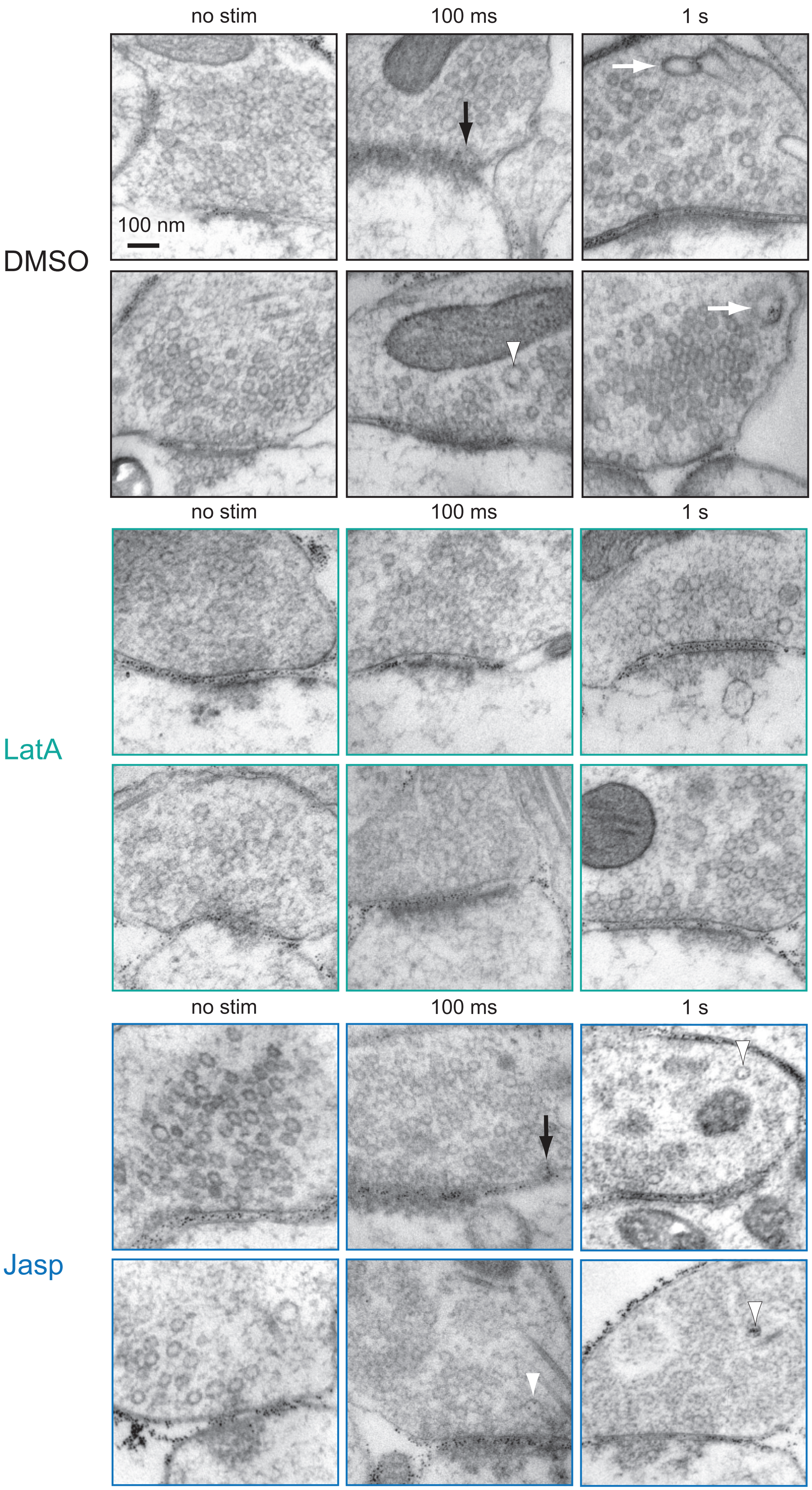

**Extended Data Figure 2.**

Additional electron micrographs showing ChetaTC-expressing wild-type neurons, treated with 0.1% DMSO, 10 μM Latrunculin A (LatA), and 100 nM Jasplakinolide. The left panel show unstimulated conditions, while the right panel show 100 ms after single stimulus (10 ms light pulse, 37 ºC, 4 mM external $Ca^{2+}$). Black arrow: endocytic pit. White arrowhead: ferritin-positive endocytic vesicle. White arrow: ferritin-positive endosomes.

### Extended Data Figure 3

a

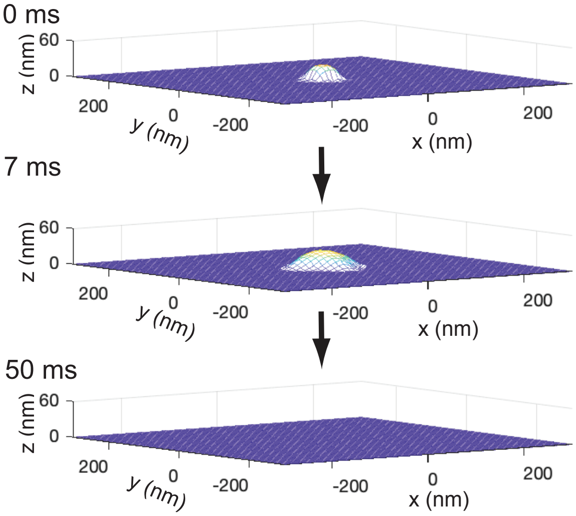

**Extended Data Figure 3.**

Snapshots from simulation, showing the evolution of membrane shape within the active zone upon a single vesicle fusion. As long as the membrane area is conserved, the fusing vesicle cannot flatten out completely. However, when the membrane area conservation was turned off at 7 ms, the exocytic pit fully collapse into the active zone.

Extended Data Figure 4

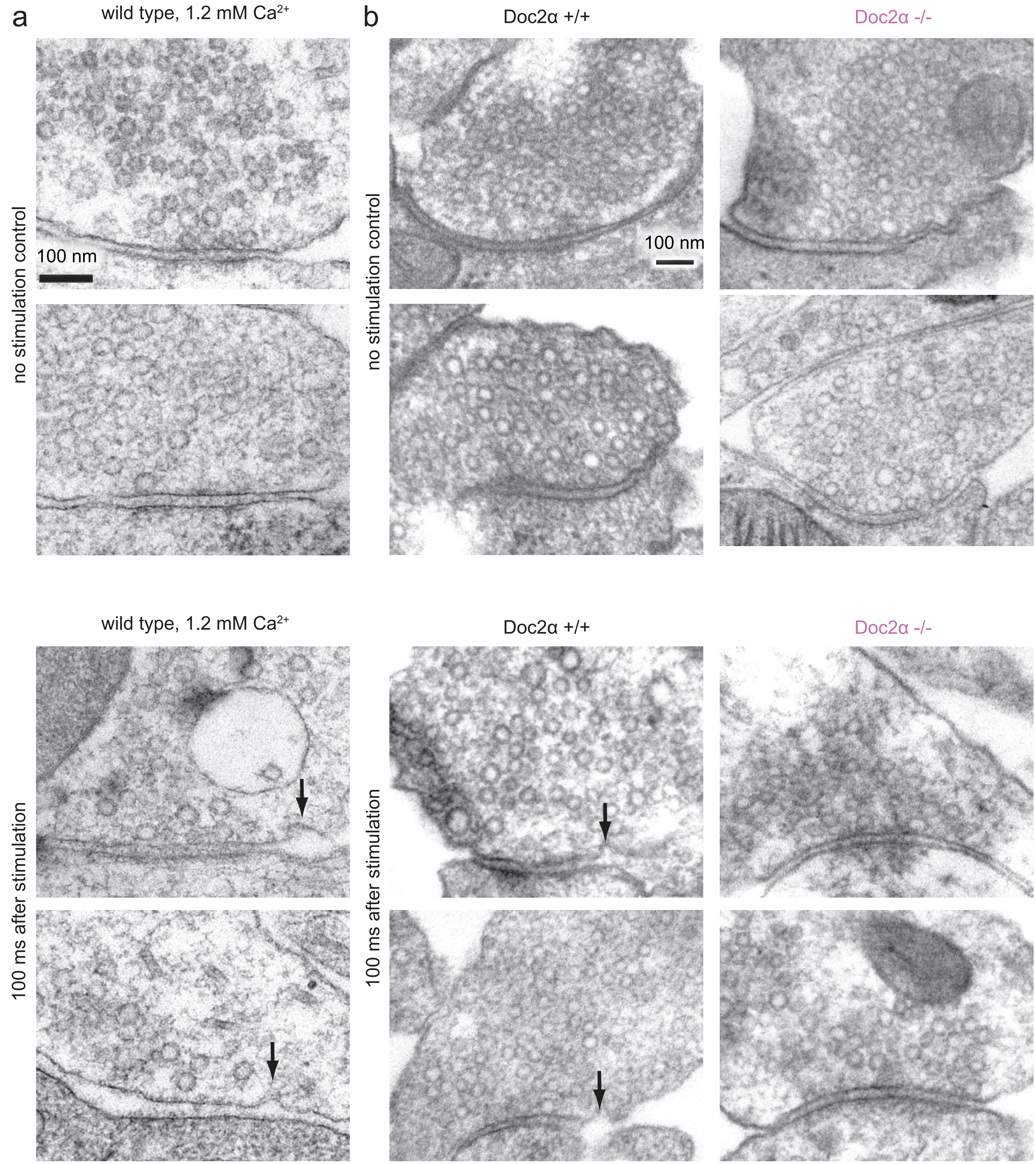

**Extended Data Figure 4.**

a. Additional example micrographs showing wild-type synapses unstimulated (left) or stimulated with electric field for 1 ms and frozen 100 ms later (right). The external calcium concentration is 1.2 mM. Black arrow: endocytic pit.

b. Additional example micrographs showing wild-type and Doc2$\alpha$ knockout synapses unstimulated or stimulated with electric field for 1 ms and frozen 100 ms later. Black arrow: endocytic pit.

Extended Data Figure 5

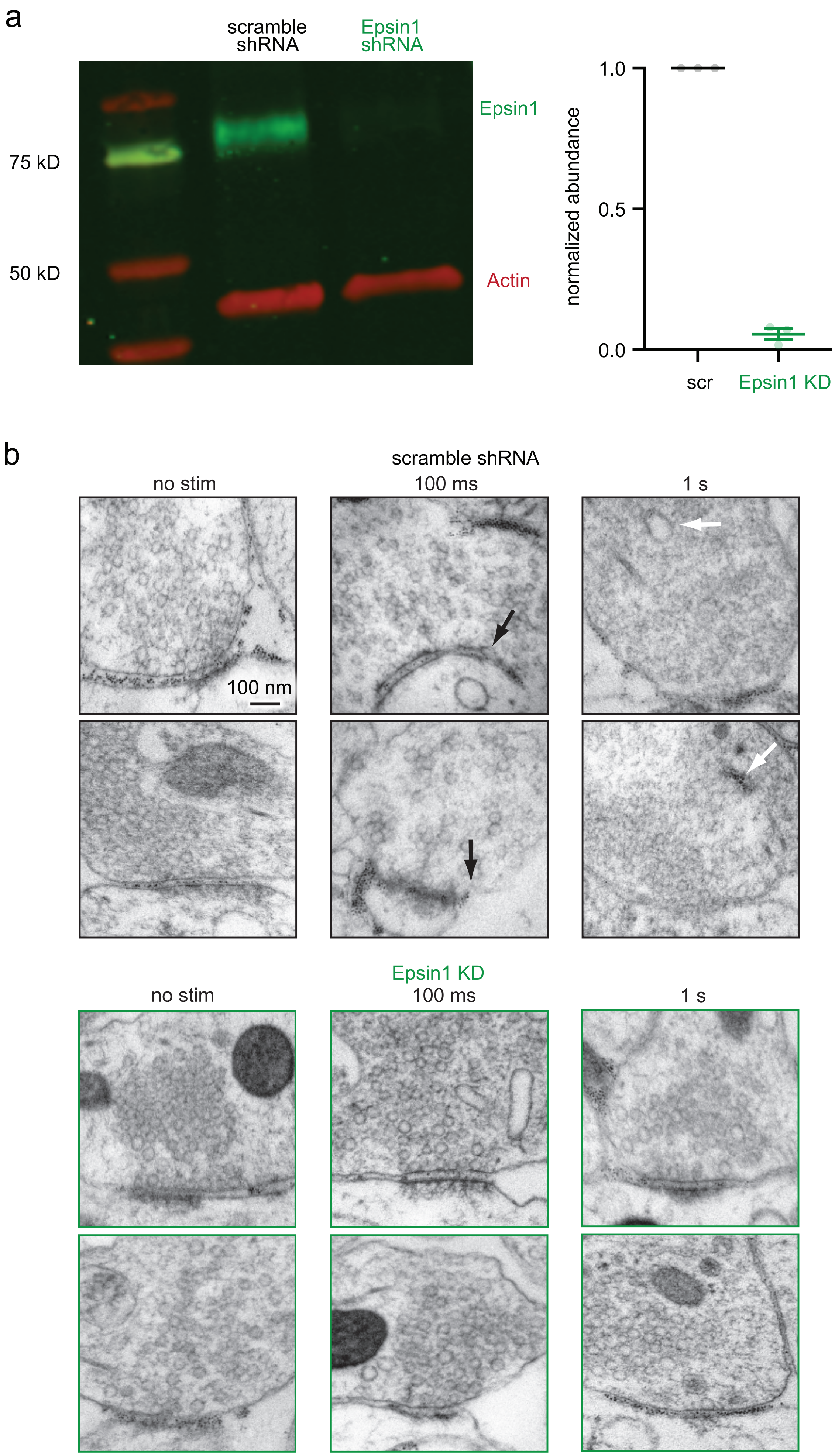

**Extended Data Figure 5.**

a. An example Western blot and a plot showing efficiency of Epsin1 knock-down (KD). In each case, Epsin1 signal intensity is normalized by the amount of actin, and then, the abundance of Epsin1 is calculated based on the normalized intensity in the scramble (scr) shRNA control. The mean and SEM are shown. Dot: one culture.

b. Additional example electron micrographs showing wild-type and Epsin1 KD synapses unstimulated or stimulated with a single electrical pulse (1 ms) and frozen 100 ms or 1 s later. Black arrow: endocytic pit. White arrow: ferritin-positive endosomes.
